# Enhanced delivery of lipid nanoparticle-based immunotherapy by modulating the tumor tissue stiffness using ultrasound-activated nanobubbles

**DOI:** 10.1101/2025.04.13.648512

**Authors:** Anubhuti Bhalotia, Pinunta Nittayacharn, Meghna Mehta, Arya Iyer, Diarmuid Hutchinson, Andrew Cheplyansky, Koki Takizawa, Abraham Nidhiry, Ramamurthy Gopalakrishnan, Theresa Kosmides, Agata A. Exner, Efstathios Karathanasis

## Abstract

Tumors often exhibit an extracellular matrix with elevated stiffness due to excessive accumulation and crosslinking of proteins, particularly collagen. This elevated stiffness acts as a physical barrier, impeding the infiltration of immune cells and the effective delivery of various immunotherapeutic agents, such as lipid nanoparticle-based RNA therapeutics. Here, we investigate the ability of ultrasound-activated nanobubbles (US-NBs) to increase the permeability and immunogenicity of tumors. Our results show that US-NBs physically remodel the tumor tissue by decreasing its stiffness by 60% five days after a single treatment. US-NB-treated tumors display randomly oriented collagen with a 5.47-fold lower deposition compared to untreated tumors. This leads to the effective delivery and widespread distribution of lipid nanoparticles (LNPs) in the tumor. When LNPs are assisted by US-NB, they have higher gene-transfection across pan-immune cells relative to LNPs alone. Notably, US-NB enables LNPs to genetically modify T cells directly in vivo. By effectively engaging both arms of the immune system, US-NB-assisted LNPs enhance the tumor immunogenicity and infiltration of cytotoxic cells by 4-fold when compared to LNPs alone. These results indicate that gentle mechanical stimulation of the tumor using US-NB offers a promising strategy to augment the delivery and efficacy of existing immunotherapies.

**Graphical Abstract:** 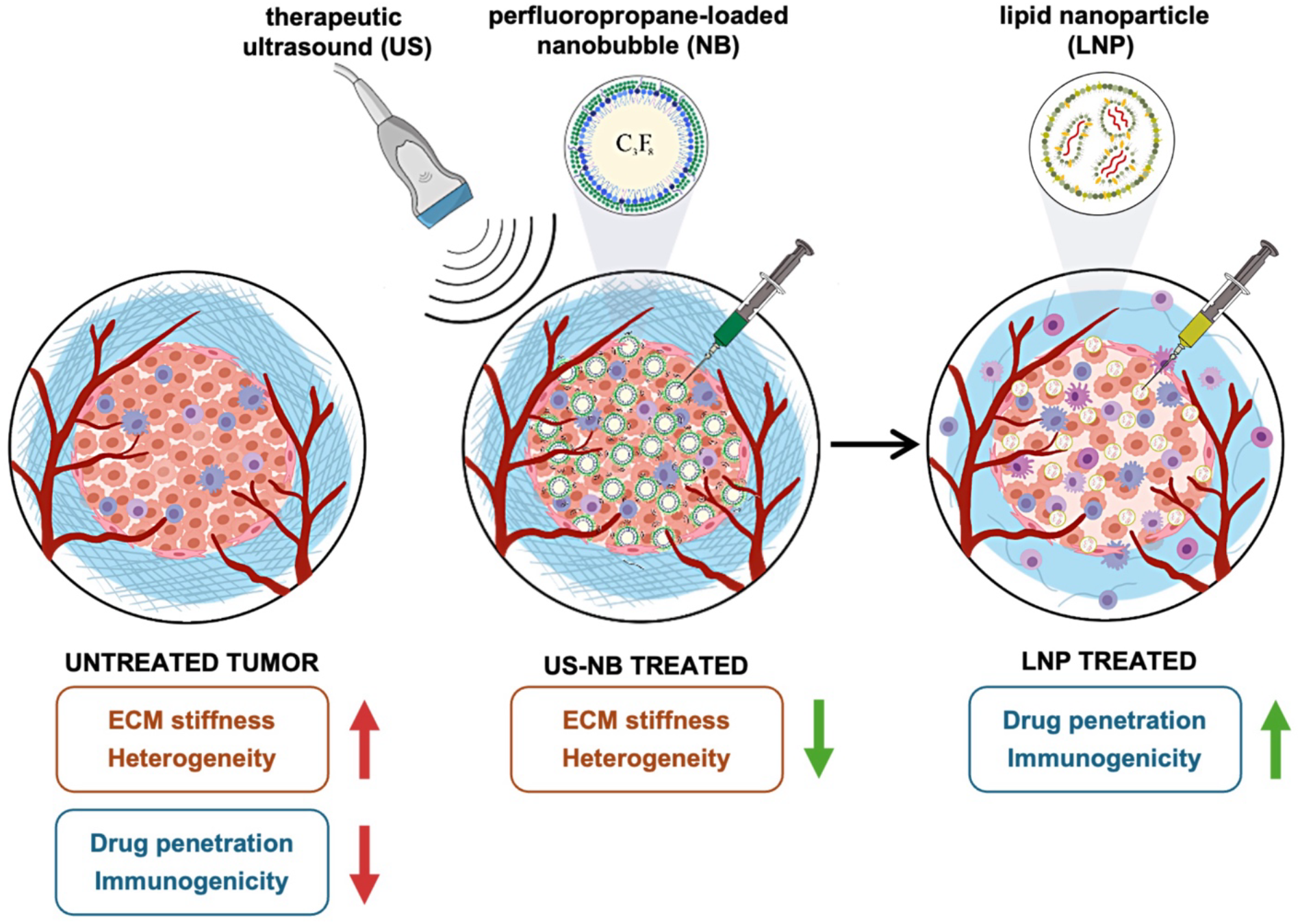

## INTRODUCTION

Cancer immunotherapies often fail because cytotoxic T lymphocytes are marginalized and excluded from cancer masses,^1,2^ due to significant challenges, including massive immunosuppression^3^ and elevated extracellular matrix (ECM) stiffness.^4,5^ The interplay between the tumor microenvironment and the ECM stiffness is established as a critical factor in regulating the malignancy and immunogenicity of growing tumors.^6,7^ Tumors remodel their ECMs, which is manifested by an excessive accumulation and crosslinking of ECM proteins, particularly collagen.^8^ First, the stiff tumor ECM compromises the proper function of both innate and adaptive immune cells.^9,10^ Further, the elevated ECM stiffness acts as a physical barrier impeding the infiltration of immune cells and the effective delivery of immunotherapeutic agents,^5,11^ including mAb for immune checkpoint blockade (ICB) adoptive cell therapy, vaccines, and lipid nanoparticle-based RNA therapeutics.

Ultrasound-activated (US) nanobubbles (NBs) are echogenic perfluoropropane gas bubbles of nanoscale diameters with a robust, resilient phospholipid shell. The deformable shell and compressible gas core of NBs allows them to penetrate deep into tissue.^12-14^ Due to these properties, NBs efficiently distribute throughout the entire tumor mass, creating widespread reservoirs of tiny gas bubbles. Stimulation of the NBs with ultrasound induces stable or inertial bubble cavitation, which, in turn, generates microstreaming and microjets, resulting in mechanical (or thermal) effects on the surrounding tissues.^15^ Our hypothesis is that gentle mechanical stresses under mild therapeutic ultrasound generated by US-NB lead to restoration of the tumor ECM elasticity. As such, acoustically stimulated NBs have the potential to remodel the tumor ECM without damaging the tissue.^16^ This can restore the efficient infiltration of immune cells and enable the proper transport of immunotherapeutic agents.

In this work, using a murine model of breast cancer, we assess the impact of US-NB on the tumor ECM and subsequent drug delivery. Using ultrasound imaging, we show that intratumorally administered NBs efficiently distribute throughout the entire tumor. The widespread distribution of NBs in the tumor produces a remarkable decrease of the tumor ECM stiffness after the application of therapeutic ultrasound as measured with ultrasound elastography and histological analysis. We use lipid nanoparticles (LNPs) to illustrate the effect of US-NB-mediated ECM remodeling on the delivery of immunotherapies. US-NB uniformly dispersed LNPs in the tumor and increased the delivery and concentration of LNPs to immune cells, remarkably even T cells. US-NB/LNPs genetically modified T cells directly in vivo, a mechanism which is typically inaccessible to delivery systems.

## RESULTS

### Nanobubbles achieve widespread distribution throughout an entire tumor

The gas core, deformable and compressible shell, and nanoscale size allow NBs to rapidly spread throughout the tumor tissues. NBs filled with C_3_F_8_ were characterized with dynamic light scattering, zeta potential, and resonant mass measurement.^17^ Fig 1a shows the hydrodynamic diameter of NBs being 283 nm with a good agreement between intensity and number-weighted size distributions. Due to the resilient shell, nearly 90% of the particles are acoustically buoyant (Fig 1b). NBs have a slightly negative surface (Fig 1c) charge (-34 mV, PBS, pH 7.4). As shown in the nonlinear contrast-enhanced ultrasound (CEUS) images (Fig 1d), NBs are able to efficiently distribute throughout the tumor, from the center (injection site) to the periphery of the tumor. Notably, NBs fill the entire tumor immediately upon intratumoral administration. To concentrate NB cavitational impact on the ECM, therapeutic ultrasound (TUS) was applied upon injection. B-mode images were taken at the center of the tumor to visualize the tumor ROI. CEUS imaging post-TUS treatment showed (last panel in Fig 1d) efficient cavitation with resulting signal comparable to baseline. To quantitatively assess NB filling and cavitation in the entire tumor, a volumetric CEUS sweep scan was conducted. The contrast signal in each frame was averaged to reveal ∼80% of each tumor slice (Fig 1e) contained NBs, indicating nearly complete NB distribution, notably even in the periphery. Similarly, contrast signal intensity post-TUS (Fig 1f) demonstrated a 68-fold loss, indicating efficient NB cavitation. The contrast signal for each frame was quantified after subtracting the baseline signal.

**Figure 1:**
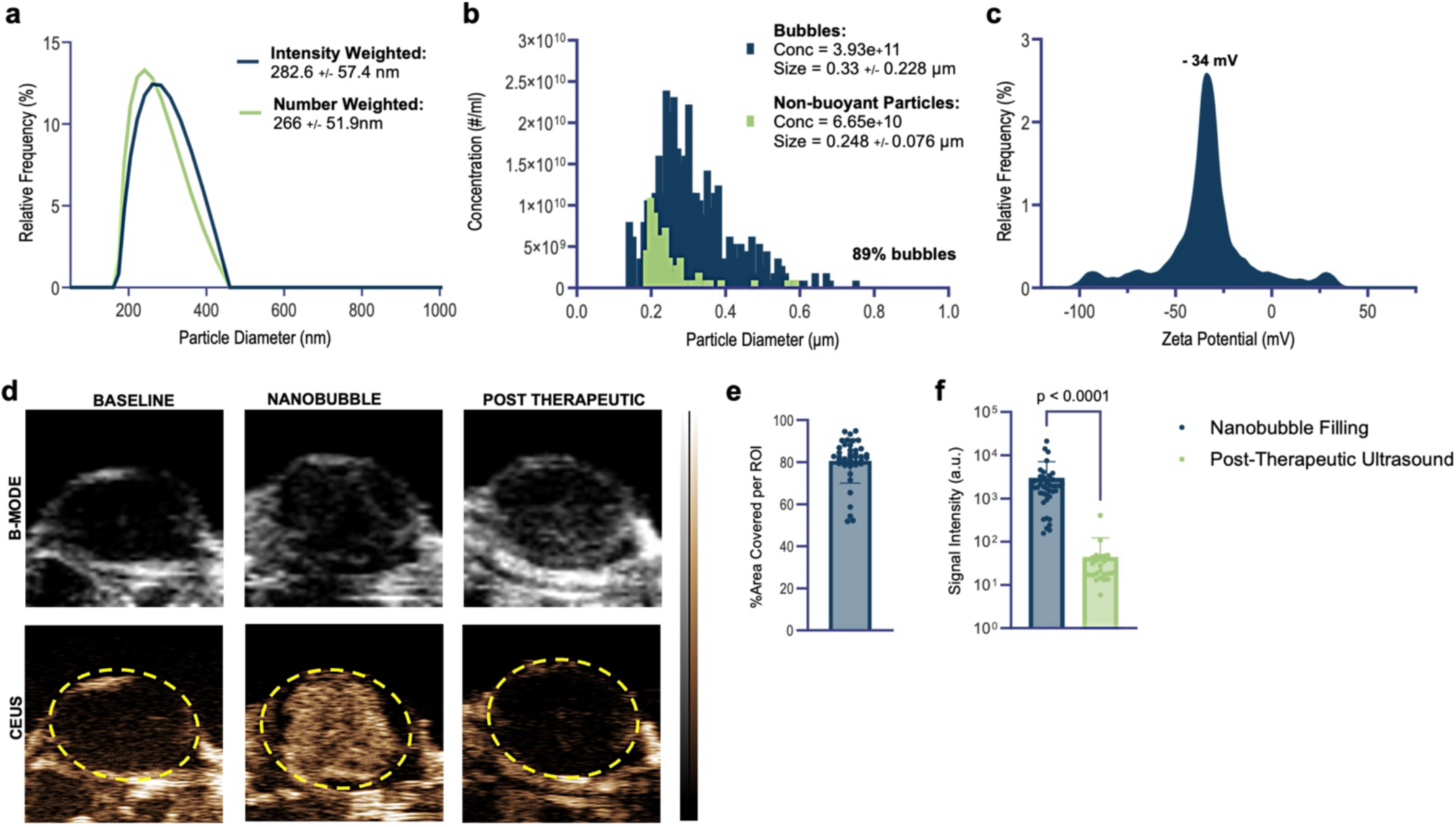
Characterization of nanobubbles and their in vivo non-linear contrast. (a) Hydrodynamic diameter of nanobubbles in PBS, both intensity and number weighted. (b) Resonant Mass Measurement of nanobubble acoustic buoyancy and concentration. (c) Zeta potential of nanobubbles in PBS at RT (d) B-Mode and Contrast Enhanced Ultrasound (CEUS) of static frames at the center of a E0771.LMB tumor at baseline, post-nanobubble injection, and post-therapeutic ultrasound. (e) Volumetric CEUS scanning used to determine %area covered per ROI for frames along the tumor length (f) Volumetric CEUS scanning used to determine non-linear contrast signal intensity for each frame along the tumor length. Statistics were performed via Student’s t test with Wilcoxon matched-pairs signed rank post-hoc.

NBs have the highest internal pressure immediately after generation.^18^ Since we want to focus the cavitational impact on the ECM, rapid filling allows for high cavitational potency. The complete spatial spread, including into the periphery of the tumor increases the likelihood of US-NB and nanobubble cavitation impacting the entire tumor. Cavitation-generated mechanical forces, such as jet streams and shock waves,^19^ can increase the permeability of the matrix, which subsequently should facilitate deeper penetration of drugs.

### Nanobubble cavitation reduces the ECM stiffness of the tumor

US-NB cavitation generates mechanical forces, such as jet streams and shock waves,^19,20^ which should increase the ECM permeability and subsequently allow for better delivery and transport of drugs in the tumor. Considering NBs retain their highest internal pressure (i.e., gas cargo) immediately after injection, the rapid distribution in the entire tumor allows us to apply TUS soon after injection to generate a significant cavitational impact on the ECM. Ultrasound shear wave elastography (SWE) was used to quantify tumor stiffness and qualitatively assess tissue heterogeneity. SWE allows non-invasive in vivo monitoring, therefore allowing repeated assessments of the same tumor over time. Prior to any measurements, the SWE probe was validated with a polyacrylamide phantom (Fig S1a). The measurements accurately returned elastic moduli for the 40 and 2kPa phantoms with high signal-to-noise ratios and homogenous elastograms. The setup for the animal studies was designed to ensure reproducible focal depth by using a standoff gel pad and acoustic coupling gel. Fig 2a shows representative US images indicating the ROIs used for analysis of the elastic modulus. The SWE measurements were averaged across 3 different imaging planes. As expected,^7^ tumor modulus in the PBS control group increased to ∼204% of the initial modulus by day 5 (Fig 2b). Immediately after US-NB, the elastic modulus of the tumor decreased to 35% of its pre-treatment value. Notably, US-NB-treated tumors sustained the lower stiffness (39% of the initial) for five days after a single US-NB application. Lower stiffness in a tumor is correlated to remodeling the ECM to a more permeable tissue structure.

**Figure 2:**
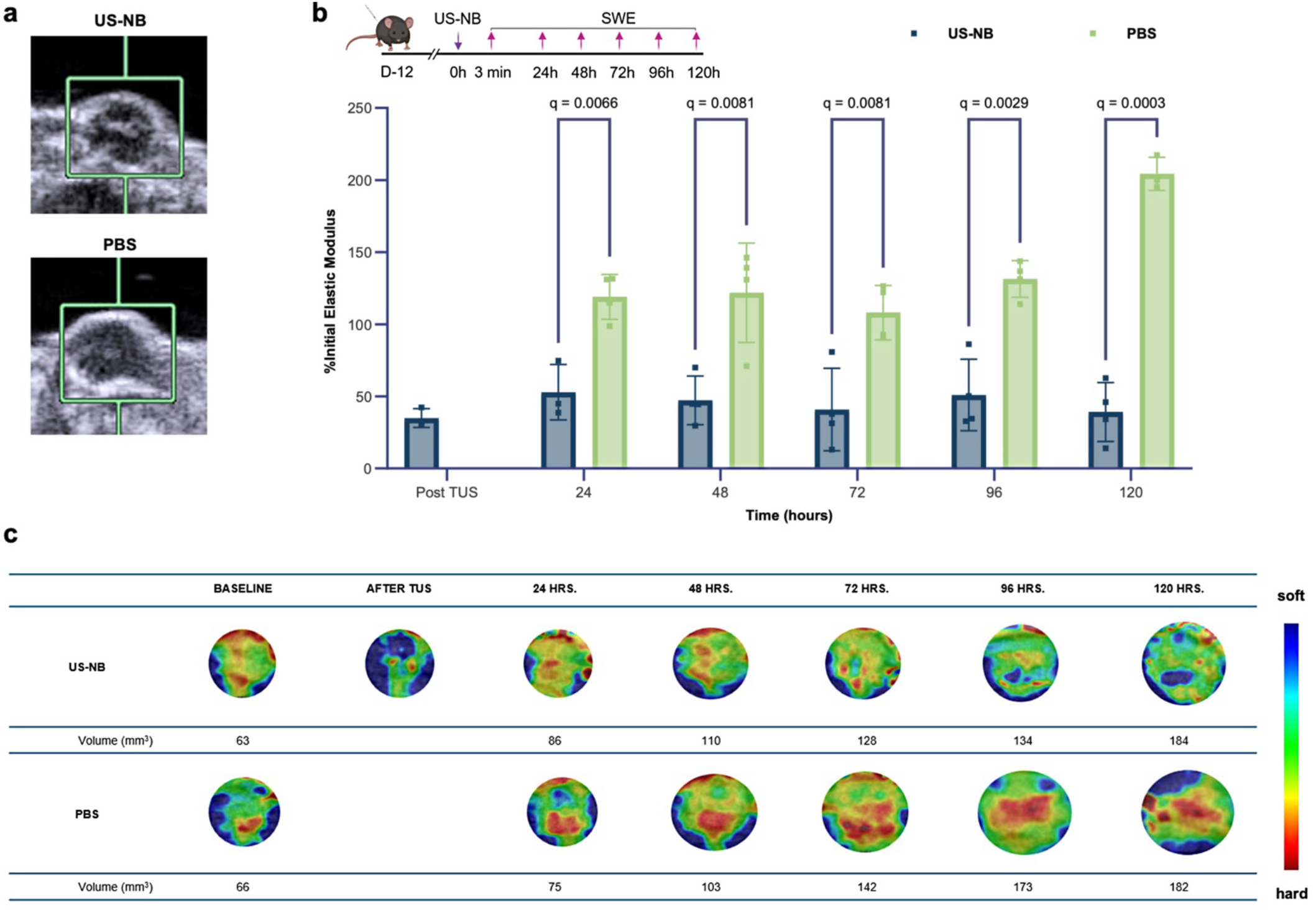
Evaluation of tumor stiffness with Ultrasound Shear Wave Elastography (SWE). (a) Representative B-mode images of E077.LMB tumors used for quantitative measurements of the elastic modulus. (b) Quantitative measurements of elastic moduli over time, every day for 5 days (n=4). Stiffness normalized to the initial values of the tumor. Only the treated group is evaluated immediately post-TUS treatment, assuming no change in the untreated group, 3 min post injection. Statistics were conducted using multiple unpaired t-test with a two-stage step-up post hoc test at 1% false discovery rate. Resulting q values are reported. (c) Qualitative elastograms are cropped to the tumor ROI and reported with the range of soft to hard tissue. Corresponding tumor volumes obtained through calipering are reported. ROIs are kept to scale.

NB cavitation reduces tumor heterogeneity. Elastograms displaying topological stiffness were used to determine spatial heterogeneity (Fig 2c). The original images are in the Supplementary Information (Fig S2). Untreated tumors injected with PBS continued to increase in stiffness heterogeneously over time. Although initially the high modulus was limited to the core, over time the stiffness increased through the entire tumor. On the other hand, US-NB treated tumors sustained decreasing tumor stiffness over time from a single treatment. Although US-NB tumor cores also began with the highest tumor stiffness, it reduced to be one of the softest regions.

It should be emphasized that US-NB is applied almost immediately after NB administration, focusing its cavitational impact on the extracellular matrix due to limited time for intracellular uptake of NBs. The physical disruption forces spatial reorganization of the ECM, reducing its stiffness and tissue heterogeneity. Stiff ECMs and heterogenous tissue are hallmarks of aggressive and metastatic cancers.^21^ Reverting tumor ECM to normal tissue structure is required to achieve infiltration and uniform distribution of drugs and immune cells to the whole tumor.

### Nanobubble cavitation remodels the extracellular matrix

Having established reduced tumor stiffness, we wanted to investigate impact of US-NBs on ECM remodeling. ECM remodeling is directly related to effective immunotherapies, which require deep penetration of therapeutic agents and their delivery to viable immune cells. In this context, we evaluated cell death, tissue remodeling and collagen degradation. E0771.LMB tumors were treated with US-NB once they were 40-60 mm^3^ and harvested 24 hours later.

Nanobubble cavitation redistributes cells but does not induce cell death. Tumors were first evaluated for cellular density by DAPI staining (Fig 3a). US-NB was the only group that showed reduced cell density. The lowered density in the US-NB group was significant when compared to both therapeutic ultrasound alone (US-PBS, p=0.0157) and untreated (PBS, p=0.002) groups. However, tumors in the US-PBS and untreated groups did not exhibit any significant differences, highlighting the need for the cavitation induced mechanical forces. Cell viability was assessed by staining for cleaved caspase 3, a protein ensuring apoptotic fate. US-NB did not induce apoptosis as denoted by the unchanged MFI of cleaved caspase 3 (Fig 3b).

**Figure 3:**
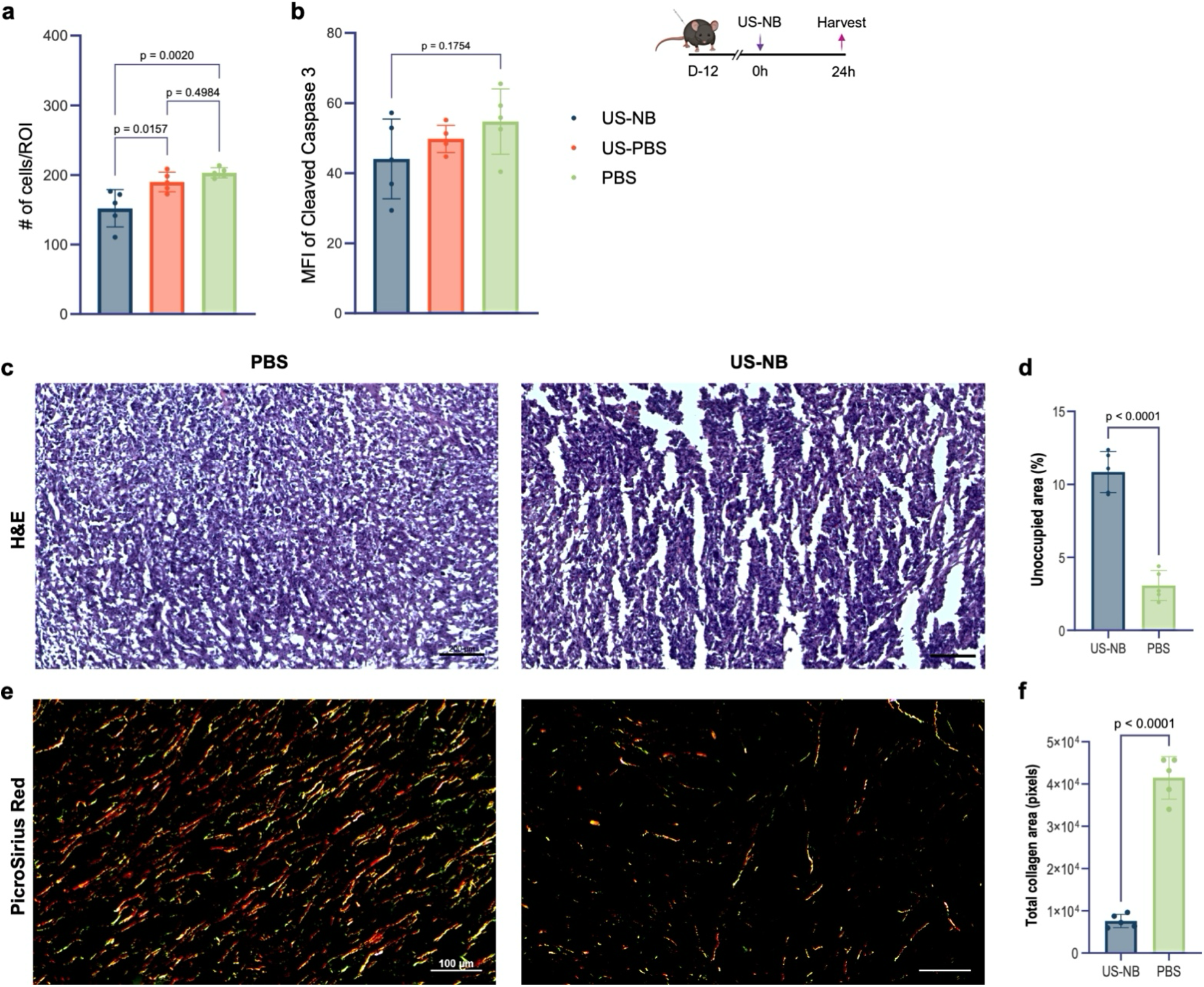
Evaluation of extracellular remodeling in the tumor. E0771.LMB tumors were treated on D12 and harvested 24 hours later for histology (n=5). (a-b) Tumor slices were stained with Cleaved caspase 3 and DAPI mounting media and imaged with a fluorescent microscope. After background subtraction, number of DAPI+ cells per ROI and MFI of cleaved caspase 3 was evaluated using MATLAB. A one-way ANOVA was conducted comparing the mean of each group to the others. (c-d) H&E staining was conducted to determine the changes in the tissue integrity. The tissue-free areas were quantified per ROI and quantified using MATLAB and student’s t-test was conducted with Welch’s correction. (e-f) Tumor slices were stained with Picrosirius Red and imaged with a polarizing microscope. Total collagen area was quantified using MATLAB and student’s t-test was conducted with Welch’s correction.

Nanobubble cavitation disrupts the structural integrity of the tissue. H&E staining revealed fragmentation of the tissue hours post-treatment (Fig 3c). US-NB generated a 3.53-fold difference in tissue-free areas compared to PBS (Fig 3d). Disrupted structural cohesivity further supports cellular reorganization as a result of nanobubble cavitation. Additionally, no necrotic regions were observed through the H&E. Remodeling tissue organization without inducing cell death, allows for enhanced tumor permeability without loss of cellular functionality.

Nanobubble cavitation remodels the extracellular matrix by collagen degradation (Fig 3c). Untreated tumors stained with Picrosirius Red revealed tumors densely populated with collagen in parallel alignment (Fig 3e). This data is expected of tumors with excessive collagen deposition. However, US-NB treated tumors had fragmented collagen, as indicated by a 5.47-fold reduction in total collagen area (Fig 3f). Additionally, US-NB interrupted collagen alignment and broke down its network. This remodeling of the tumor ECM^8^ can weaken the physical barrier to drug delivery and have significant benefits for immunotherapies.

### Nanobubble cavitation can directly deliver nanoparticles to T cells and tumoral periphery

We next aimed to determine the impact of US-NBs on drug delivery and cancer immunotherapy. For the purposes of this paper, we assessed drug delivery through lipid nanoparticles (LNPs) carrying immune checkpoint targeted siRNA.^22^ The LNPs are 50-80 nm in hydrodynamic diameter (Fig S3a) with a neutral surface charge (Fig S3b). Additionally, the LNPs have high encapsulation efficiency (>90%) of the siRNA (Fig S3c).

US-NB enhances the distribution of LNPs throughout the entire tumor (Fig 4a). To evaluate the spatial distribution, fluorescent LNPs were injected in the center of tumors, which were harvested 24 hours post-treatment for histological analysis. Without US-NB, LNPs remained in the tumor core and were not able to spread towards the periphery. However, US-NB caused a substantial increase of the distribution of LNPs (Fig 4b). Comparing the LNP accumulation in the center to the periphery of the tumor, US-NB resulted in a 4-fold increase in the accumulation in the tumor’s periphery when compared to LNPs alone (Fig 4c).

**Figure 4:**
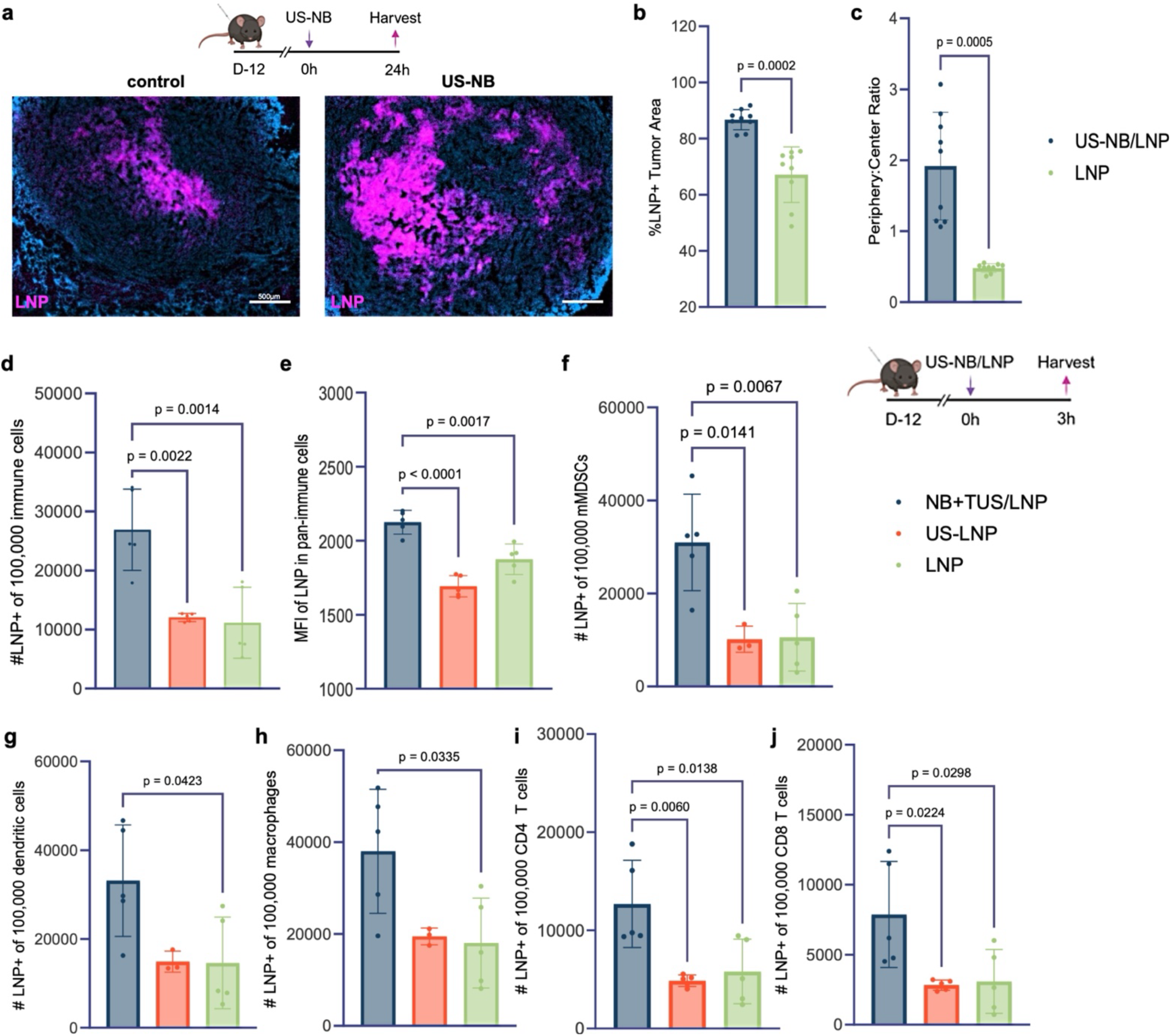
Spatial and cellular evaluation of LNPs in vivo. (a) Spatial spread of fluorescent nanoparticles (DiR+) analyzed in an E0771.LMB tumor 24 hours after treatment. Imaged with the Zeiss Axio Z1 microscope at 5x (n=3). (b) %Area of the tumor which is LNP+ across all pixels in the tumor that are DAPI+. Statistics conducted using student’s t-test with Welch’s correction. (c) LNP+ pixels reported as a ratio of the periphery to center using MATLAB segmenting. Three slices were taken per tumor. (d-e) Within the immune population, the proportion of immune cells (CD45+) that were nanoparticle positive and their mean fluorescent intensity of that expression was assessed for US-NB/LNP, US-PBS-LNP, LNP. (c) Macrophages (CD11b+F4/80+MHC2+), (d) dendritic cells (CD11b+CD11c+), (e) mMDSC (CD11b+Ly6G-Ly6Chi), (f) CD4 T cells (CD3e+CD4+) and CD8 T cells (CD3e+CD8+) cells were similarly quantified. Experiments were conducted with n=5 mice per group. Injection volumes were kept consistent. Statistics were performed using a one-way ANOVA, comparing the mean of each group with the others.

US-NB increases the uptake of LNPs by immune cells. Nanobubble cavitation enhanced LNP internalization by tumor-resident immune cells. E0771.LMB tumors were harvested 3 hours after treatment by US-NB and LNP, US and LNP, or LNPs alone. All treatment volumes and number of injections were equalized, supplemented with PBS when needed. US-NB caused the delivery of LNPs to at least 2.3-fold more immune cells when compared to US and LNPs without US (Fig 4d). Furthermore, LNP accumulation per immune cell was significantly higher for US-NB relative to its controls (Fig 4e).

US-NB enhances the delivery of LNPs to T cells and innate immune cells key to an antigen-mediated response.^23^ US-NB significantly increased LNP delivery to mMDSCs (Fig 4f), dendritic cells (Fig 4i), and macrophages (Fig 4h) by an at least 2-fold higher uptake when compared to controls. Considering that LNP delivery to T cells is challenging due to their poor endocytic mechanisms, US-NB-LNPs are directly internalized by both CD4 (Fig 4i) and CD8 T cells (Fig 4j). US-NB resulted in LNPs being internalized by ∼3-fold more T cells compared to the control conditions. Overall, US-NB improves the spatial distribution of LNPs in the tumor and their cellular uptake by different immune cell subsets, including both the innate and adaptive immune cells.

### Nanobubble cavitation enhances genetic transfection

Since the overall delivery of LNPs to the tumor is enhanced, we next wanted to determine US-NB effects on the LNP-mediated transfection efficiency. To limit any biological interference from nanobubble cavitation, we evaluated transfection of an exogenous gene (eGFP mRNA). LNPs were made using syringe mixing and characterized to reveal a hydrodynamic diameter of ∼69 nm.

US-NB augments LNP-mediated transfection and gene expression levels for pan-immune cells. Tumor volumes were compared to an untreated group, to demonstrate no changes in growth rates. E0771.LMB tumors were treated with eGFP mRNA loaded LNPs and harvested 24 hours later. GFP expression was evaluated on a single cell level using flow cytometry. US-NB increased the transfection efficacy of LNPs across pan-immune (CD45+) cells by 1.4-fold (Fig 5a). Moreover, US-NB resulted in higher MFI (Fig 5b), indicating an increase in the GFP expression. Increased transfection was also observed across activated macrophages, dendritic cells and mMDSCs by ∼1.5-fold (Fig 5c-e). Overall, US-NB caused an increase in the number of cells being transfected as well as the levels of gene expression.

**Figure 5:**
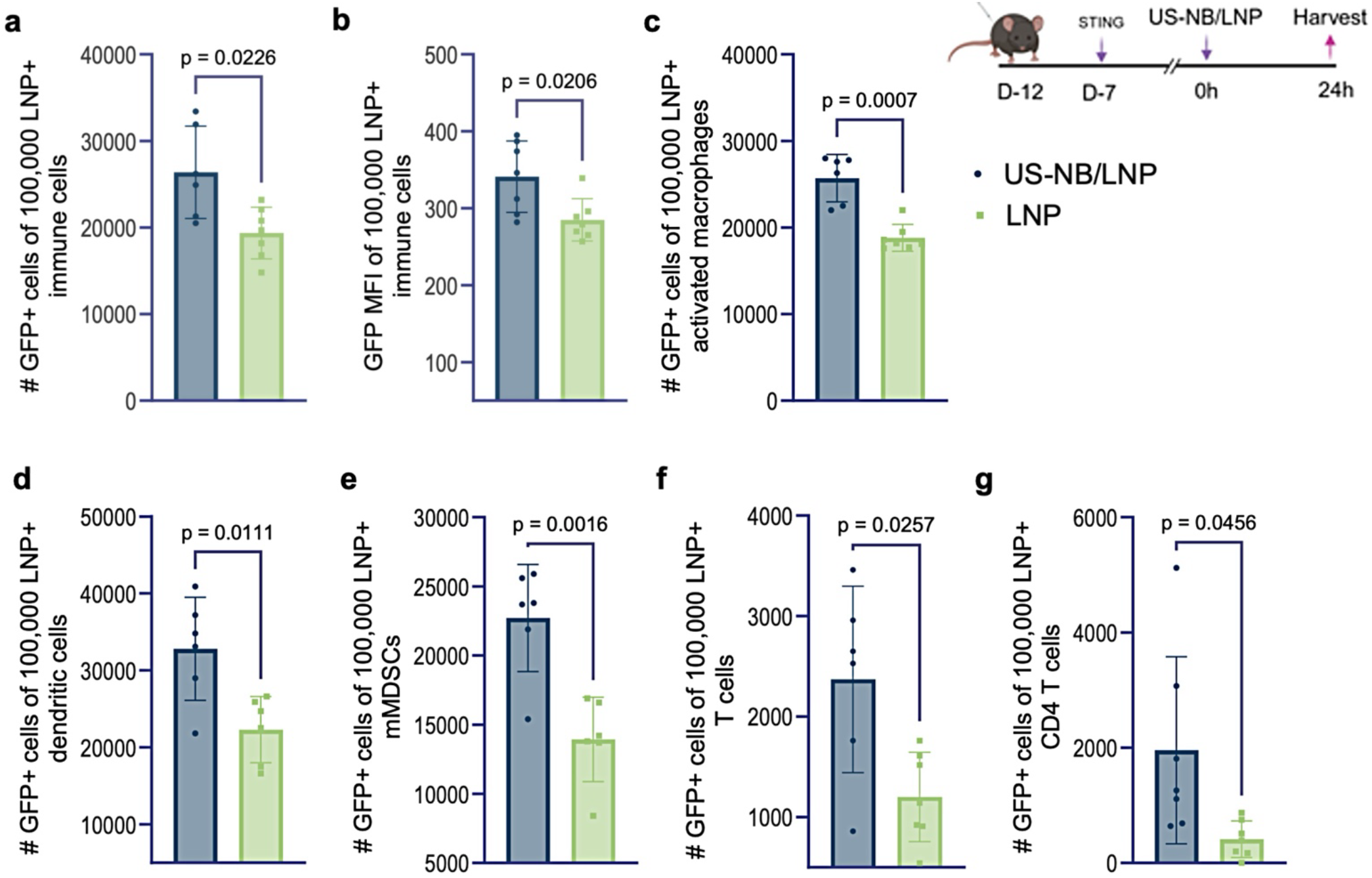
Improved transfection efficiency of US-NB-LNPs in tumor-resident immune cells. Mice were inoculated with E0771.LMB tumors and were subcutaneously injected with STING agonist 5 days post-inoculation to activate the tumor-resident innate immune cells. E0771.LMB tumors were treated when they were 40-60 mm^3^ with GFP mRNA-carrying fluorescent lipid nanoparticles. Tumors were harvested and assessed at a single cell level by flow cytometry. (a-b) Within the population that was nanoparticle positive, the proportion of immune cells (CD45+) expressing GFP and the mean fluorescent intensity of that expression was compared. (c) Activated macrophages (CD11b+F4/80+MHC2+), (d) dendritic cells (CD11b+CD11c+), (e) mMDSC (CD11b+Ly6G-Ly6Chi), (f) T cells (CD3e+) and CD4 T cells (CD3e+CD4+) cells were similarly quantified. Experiments were conducted with n=6 mice per group. Injection volumes were kept consistent. Statistics were performed using an unpaired student’s test with Welch’s correction.

US-NB increases the LNP-mediated transfection of T cells. Remarkably, upon nanobubble cavitation, double the number of LNP+ T cells expressed GFP (Fig 5f) compared to LNP delivery without US. Notably, the CD4 subset of T cells exhibited a 6-fold increase in transfection (Fig 5g). Typically, CD4+ T cells in the tumor are primarily regulatory or exhausted and therefore are a desirable target for LNP-mediated therapy. US-NB improves the LNP-mediated transfection and transgene expression in pan-immune cells. For immunotherapies to succeed, they need to engage both the innate and adaptive immune system.^24^ US-NB/LNPs can increase transfection across all subtypes key to tumor reprogramming without requiring specific targeting. Most remarkably, US-NB can effectively direct LNPs to the adaptive immune system and directly transfect them. Reprogramming both resident immune subsets has the potential for an accelerated, antigen-specific immune response.^23^

### Nanobubble assisted immunotherapy significantly improves tumor immunogenicity

We then investigated whether the combination of US-NB and LNPs can produce effective reprogramming of the tumor immune microenvironment. We assessed immune signaling, which is crucial in generating a robust anti-tumor response. Through effective immune signaling, both the innate and adaptive immune cells can be engaged, limiting tumor immune evasion. For the purposes of this study, the LNPs were loaded with the clinically validated CTLA-4 and PD-1 specific siRNA. E0771.LMB tumors were treated three times, with each treatment being 3 days apart. Controls included US-NB alone, LNP alone, and PBS. Tumors and plasma were harvested 24 hours later and assessed with a multiplex bead assay.

The combination of US-NB and LNPs elevates chemokines key to immune infiltration. The tumor and blood showed increased levels of CXCL10 (Fig6 a-b) and CCL2 (Fig6 c-d). For example, US-NB/LNP resulted in at least a ∼2-fold enhancement of CXCL10 in the tumor relative to the controls (Fig 6a). CXCL10 signaling recruits CD8+ T cells and NK cells into the tumor, generating a cytotoxic response. Similarly, US-NB/LNP generated a 2-fold increase in the CCL2 levels in the tumor compared to controls (Fig 6c). CCL2 is correlated with macrophage recruitment and neutrophil-mediated responses.

**Figure 6:**
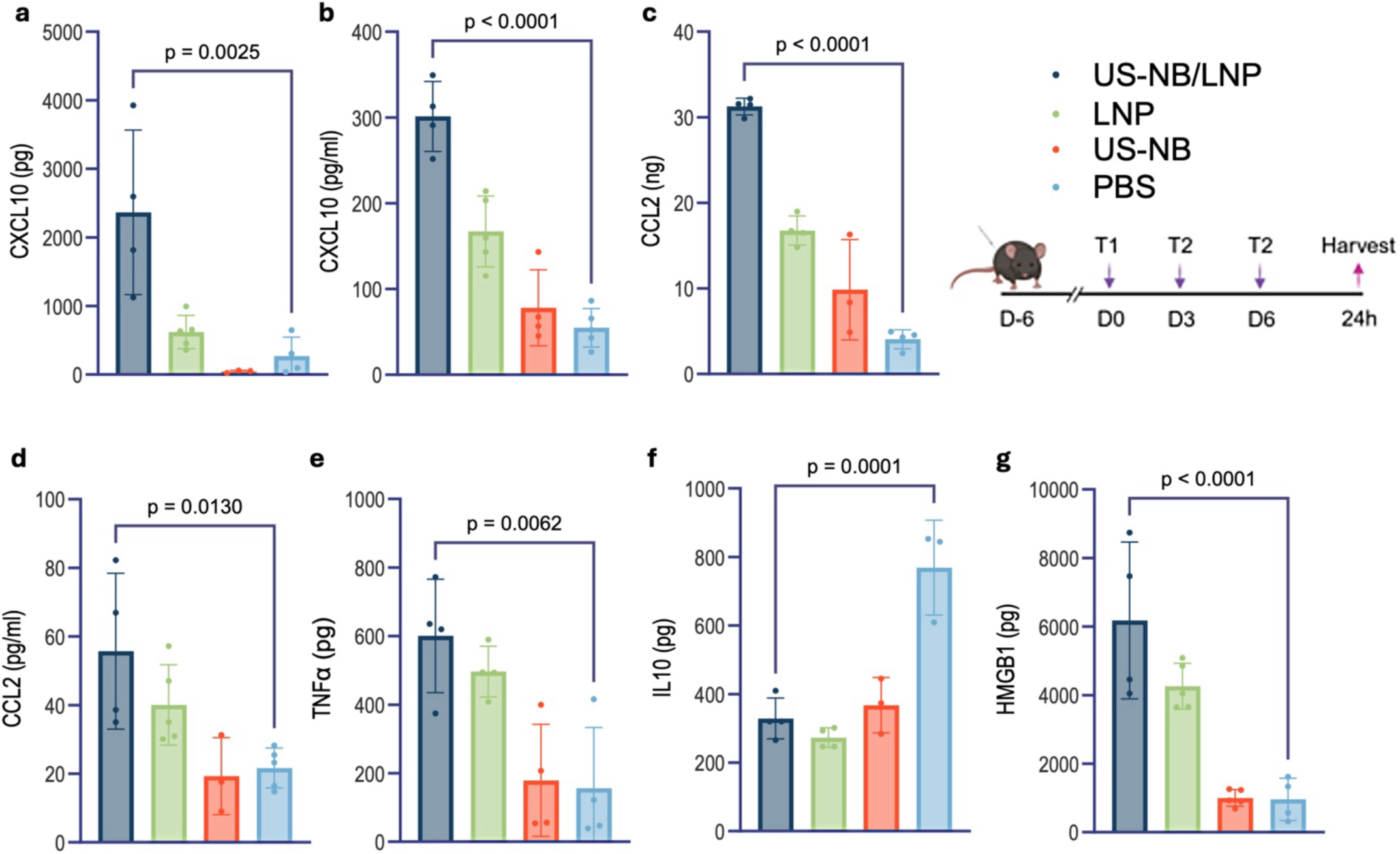
Immune signaling for combination LNPs to assess long-term changes. E0771.LMB tumors were treated on day 6,9,12 and harvested 24 hours after the third treatment. Signaling molecules were quantified using a multiplex bead assay (n=4). (a-d) Chemokines were measured in tumor homogenates and plasma for US-NB-NP, LNP, US-NB and PBS. (e-f) Similarly, pro (TNFα) and anti-inflammatory (IL-10) cytokines were measured across groups in the tumor. (g) HMGB1 in the tumor was quantified by an ELISA as a measure of ICD. All concentrations were multiplied by the homogenization volume. All injections volumes across groups were kept constant. Statistics were carried out using one-way ANOVA comparing the mean of each group to the others.

The combination of US-NB and LNPs reprograms the tumor to be pro-inflammatory. To evaluate reprogramming, TNFα, IFNγ and IL-10 levels in the tumor were analyzed. US-NB/LNP upregulated IFNγ (Fig S4) and TNFα (Fig 6e) ∼3.9 and ∼2.6-fold higher than the untreated, respectively. This is further supported by the 2.3-fold downregulation of IL-10 in the case of US-NB/LNP compared to the untreated (Fig 6f). To further investigate the immune response, we examined HMGB1 expression (Fig 6g). US-NB/LNP caused a 6.5-fold increase in HMGB1 levels when compared to untreated, indicating an increase in tumor immunogenicity that is beneficial to an antigen-mediated response.

US-NB/LNPs create an immunogenic tumor that engages both innate and adaptive immune cells. US-NB assisted CTLA-4 and PD-1 checkpoint silencing drives immune recruitment and limits immune tolerance. The generated signaling profile supports a positive prognosis and durable clinical responses. Collectively indicating that US-NB augments drug efficacy long term.

## DISCUSSION

In this study, we demonstrate that ultrasound-mediated nanobubble cavitation enhances drug delivery of lipid nanoparticles to immune cells in tumors. The improved delivery stems from the US-NB-induced remodeling of the tumor ECM. Specifically, US-NB reduces the overall tumor ECM stiffness as well as the topological heterogeneity of ECM stiffness, improving the infiltration and transport of therapeutics (and immune cells) in the entire tumor. Furthermore, data suggest that the generation of cavitational forces in the tumor microenvironment sensitizes resident immune cells, leading to improved uptake by various immune cell subsets. Most remarkably, US-NB enabled LNPs to be internalized by T cells, leading to their increased gene transfection in vivo.

Elevated ECM stiffness appears to play a key role by supporting tumor growth via various mechanisms. The increased collagen deposition, ECM stiffness, and its heterogeneity support exponential growth partially due to biomechanical forces that strongly regulate the behavior of tumor cells and function of immune cells.^25^ Equally important is that the stiff tumor ECM acts as a physical barrier impeding the infiltration of immune cells and effective drug delivery. To address this challenge, typical ultrasound interventions employ systemic administration of microbubbles, which have shown minimal tissue penetration resulting in the cavitation effects being primarily localized within the vascular and near-perivascular regions of tumors. In this study, we elected to utilize intratumoral injection to illustrate the ability of NBs to spread in tissues due to their nanoscale sizes, deformable shell and compressible gas core. Due to NBs quick penetration of the tumor, nanobubble cavitation results in a significant reduction of ECM stiffness, collagen cross-linking and tissue heterogeneity.

US-NB and the reduced ECM stiffness allowed for improved delivery of LNPs. In the absence of US-NB, the intratumoral administration of LNPs resulted in limited distribution in the tumor, with the majority of the LNPs being localized in close proximity to their injection site.^26^ This is primarily a result of high interstitial pressures in tumors causing nanoparticles to rely on diffusive distribution.^27^ Considering the tumor periphery is a region of very high cell densities and pressures, it is not surprising the transport of the LNPs from the center (injection site) to the periphery of the tumor was limited. In addition to US-NB’s impact on the broader distribution of LNPs in the tumor, the localized shock waves and microjets can sensitize cells to delivery.^28^ The sensitization of cells is typically attributed to the cavitation-induced activation of endocytic pathways. In addition to LNPs being able to disperse throughout the tumor, US-NB facilitates the internalization of LNPs by more immune cells as well as the internalization of a higher number of LNPs by each immune cell.

In this work, we focused on LNPs for immune-checkpoint silencing, US-NB can benefit many cancer therapies such as CAR-T cells, antibody treatments and cancer vaccines. Although LNPs are an increasingly popular choice for gene delivery, only a small portion of the gene cargo is intactly delivered to the cytosol.^29^ This is even more problematic in the case of LNP delivery to T cells. To direct nanoparticles to T cells, targeting schemes are employed by decorating the particles with targeting ligands, often antibodies. US-NB offers a convenient alternative by improving the transfection of pan-immune cells, exceptionally even T cells, without the need for additional modification of a nanoparticle. Thus, US-NB is agnostic to the type of LNP formulation or its cargo and can be a versatile companion for various applications of direct in vivo gene modification of antigen-presenting cells, T cells, or both innate and adaptive immune cells.

In conclusion, nanobubble cavitation remodels the ECM, resulting in enhanced delivery and efficacy of immunotherapy. While most efforts focus on optimizing the delivery system to achieve improved delivery to tumors, US-NB seeks to optimize the tumor for delivery. US-NB seems to be suitable for solid tumors, particularly those with a high stromal content. Further research development is required to understand its impact on more pliable tumors, such as melanoma. Overall, US-NB can become a drug-agnostic immunomodulatory intervention offering a wide applicability for patients with different treatment requirements.

## METHODS

### Animal studies

All animal studies were performed under an approved protocol (protocol 2016-0115) by the Institutional Animal Care and Use Committee of Case Western Reserve University. E0771.LMB (ATCC, Manassas, VA, USA) were cultured in DMEM supplemented with 10%FBS and 1%PS (Gibco, Evansville, IN, <u>USA</u>). 6-8 week C57BL/6J (Jackson Labs, Bar Harbor, ME, USA) were intradermally inoculated on the right flank with 250,000 tumor cells in 30μl of DMEM. Mice were monitored daily for their weight and tumor volumes (length x width^2^ x 0.5), where length is the longest axis, and the width is perpendicular. In the studies evaluating the effect of US-NB on tissue stiffness (Fig 1-4), tumors were treated on day 9 with nanobubble cavitation. In the mRNA transfection studies (Fig 5), tumors were first subcutaneously injected with 10 μg of STING agonist on day 3 and then treated with US-NB and LNPs on day 9. In the immunotherapeutic studies (Fig 6), tumors were treated every three days for three treatments starting on day 6.

### Synthesis of nanobubbles

NBs were formulated as previously described.^4^ Briefly, NBs were prepared by dissolving DBPC, DPPA, DPPE (Avanti, Birningham, AL, USA), and mPEG-DSPE (Laysan,Arab, AL, USA) in propylene glycol (PG). A mixture of glycerol and PBS was then added to the lipid solution. Next, 1 mL of the lipid solution was aliquoted into a 3-mL vial, and the air inside was removed and replaced with C_3_F_8_ gas. To activate NBs, the vial was shaken using a VialMix shaker (Bristol-Myers Squibb Medical Imaging, Inc., N. Billerica, MA, USA) for 45 s to induce bubble self-assembly. NBs were then isolated by differential centrifugation at 50 rcf for 5 min with the vial inverted. Fluorophore-conjugated NBs were prepared using the same method, with DSPE-PEG-Cy5 (0.1 mg/mL in chloroform) included in the formulation. The chloroform was evaporated in an 80°C water bath before lipid solution preparation.

### Characterization of nanobubbles

The size distribution and concentration of buoyant particles (bubbles) and non-buoyant particles (lipid aggregations, micelles) were analyzed using resonant mass measurement (RMM) (Archimedes, Malvern Panalytical Inc., Westborough, MA, USA) with a calibrated nanosensor (100 nm - 2 μm). The sensors were pre-calibrated using NIST-traceable 565 nm polystyrene bead standards (ThermoFisher 4010S, Waltham, MA, USA).^5,6^ NBs were diluted 1:1000 in PBS, and at least 500 particles were measured per trial (n=3). The average particle size (buoyant and non-buoyant) and aggregation were further assessed using dynamic light scattering (DLS) (Litesizer™ 500, Anton Paar, Ashland, VA, USA). NBs were diluted 1:1000 in PBS and measured for intensity- and number-weighted size distributions. NBs underwent quality control by assessing their in vitro acoustic properties, including initial contrast enhancement and stability under ultrasound, using a tissue-mimicking agarose phantom.^5^ Nonlinear contrast images were continuously acquired using an ultrasound scanner (Vevo 2100, VisualSonics, New York, NY, USA) with nonlinear contrast-enhanced ultrasound at 18 MHz, 4% power, and 1 frame per second over 5 minutes. Raw data were recorded and analyzed using ImageJ software (freeware available from NIH). The region of interest (ROI) was defined, and total intensity was quantified using the Time Series Analyzer V3 plugin. From the data, initial signal enhancement, signal decay over time, and the percentage of remaining signal at 8 minutes were determined. The experiment was conducted in triplicate.

### Therapeutic ultrasound

NB cavitation studies were carried out in the murine E0771.LMB model when the tumor size was ∼50 mm^3^. NBs were diluted 1:10 in PBS, and ∼7×10^7^ NBs were intratumorally administered using a 29G½ insulin syringe (Exel International, Salaberry-de-Valleyfield, Quebec J6T 0E3, Canada). Therapeutic ultrasound (TUS) was applied on the tumor using an unfocused transducer with a 1 cm^2^ effective radiating area (Sonicator 740, Mettler, Anaheim, CA, USA) at 3.3 MHz, 2.2 W/cm^2^, and a 50% duty cycle for 1 min. TUS parameters included a pulse repetition frequency (PRF) of 100 Hz, a pulse length of 10 ms, and an estimated peak negative pressure (PNP) amplitude of 0.25 MPa.

### Volumetric non-linear contrast-enhanced ultrasound imaging

Three-dimensional tumor imaging was performed using a preclinical ultrasound imaging system (Vevo F2, FUJIFILM VisualSonics, New York, NY, USA) equipped with a UHF29X transducer (15–29 MHz bandwidth). Imaging was conducted in dual B-mode and 3D nonlinear contrast (NLC) mode, with an NLC transmit frequency of 17.5 MHz (10% power, 0 dB contrast gain, 0 dB gain, 50 dB dynamic range, 5 frames per second, 0.2 mm step size). The focus was aligned with the tumor center, and imaging began immediately after NB injection. The region of interest (ROI) was defined, and the contrast-enhanced signal was quantified using MATLAB (MathWorks, Natick, MA, USA).

### US-Shear wave elastography (SWE)

Tumor elasticity was measured using an ultrasound scanner (Siemens Acuson S2000, Siemens Healthineers, Munich, Germany) equipped with a 9L4 linear array probe (4-9 MHz bandwidth). To ensure a consistent scanning setup, the probe-to-tumor distance was maintained using a standoff gel pad (Aquaflex, Santa Ana, CA, USA) and a uniform layer of acoustic coupling gel. Scans were performed perpendicular to the tumor with a fixed ROI depth, following the manufacturer’s recommended settings for SWE. Tumor-bearing mice received therapeutic ultrasound treatment and saline intratumoral injections on day 0 and were monitored every day for 5 days. The nanobubble cavitated group was additionally assessed immediately after TUS and at 3 hours post-treatment. To validate SWE measurements, polyacrylamide phantoms with specific moduli were utilized. The preparation process followed a previously reported method.^7^ In summary, phantom solutions were prepared using 40% acrylamide at concentrations ranging from 6.4 to 26.5 wt.% (Supplementary Table 1). Cross-linking was initiated through free radical polymerization, facilitated by TEMED and APS, with polymerization occurring at room temperature within designated containers. Each phantom contained 0.095 wt.% bis-acrylamide as a cross-linking agent. After polymerization, the phantoms were stored in PBS to maintain stability.

### Histological analysis

E0771.LMB-bearing mice were given one treatment of therapeutic ultrasound treatment with nanobubbles and PBS, or only PBS and tumors were harvested after 24 hours. Tumors were washed in PBS and then placed in 30% sucrose for 48 hours followed by 4% PFA at 4°C. Tumors were then washed and placed in OCT for 72 hours at -80°C before being cryo-sliced at 12 μm. Cell death IF staining used a polyclonal Ab specific for cleaved caspase 3 (Life Technologies, CA, USA) and Alexa Fluor 488 Antibody (Life Technologies, CA, USA) and DAPI mounting media (Abcam, Waltham, MA, USA). H&E and picrosirius staining for collagen was conducted by the research core. IF slides and H&E brightfield imaging was conducted on the Zeiss Axio Z1 (Carl Zeiss Microscopy, White Plains, NY, USA). Picro Sirius staining was imaged with the Zeiss Axio Z1scanner.

### Synthesis and characterization of lipid nanoparticles

Lipid nanoparticles (LNPs) loaded with siRNA were synthesized as previously described .^8^ DLin-MC3 (Cayman Chemical), cholesterol (ovine) (Avanti Lipids), Distearoylphosphatidylcholine (DSPC) (Avanti Lipids), 1,2-dimyristoyl- rac- glycero- 3- methoxypo lyethylene glycol-2000 (DMG-PEG2000) (Avanti Lipids) and 1,2-Distearoyl- sn- glycero- 3- phosphoethanolamine- mPEG (DSPE-mPEG2000) (Laysan Bio) were mixed in a solution of chloroform at a ratio of 50:38:10.5:1.4:0.1. Lipid solutions were evaporated at room temperature and the resulting films were hydrated with siRNA in nuclease-free acetated buffer in a 1:3 ratio (v/v) and mixed rapidly. The lipid-drug solution was then probe sonicated at 20% PW at 30s on, 10s off for 5 min. The solution was encapsulated in 300 kDa membranes and dialyzed against 1X nuclear free-PBS for 16-18 hours. LNPs loaded with mRNA were made with an identical lipid solution but with the ionizable cationic lipid switched from D-Lin-MC3 to SM102 of the same molar ratio. EGFP mRNA (TriLink Biotechnologies) was reconstituted in nuclease-free citrate buffer and the lipid solution was mixed with the mRNA at a 1:3 ratio (v/v) through syringe mixing. The solution was encapsulated in 20 kDa membranes and dialyzed against 1X nuclear free-PBS for 16-18 hours. The resulting nanoparticles were concentrated by centrifugation at 3000 rpm in an Amicon filter unit. When fluorescently tagged, 0.2-0.5% DiR’ was added to the lipid solution, taking away the mol% from cholesterol. Nanoparticles were characterized for their hydrodynamic size using dynamic light scattering (DLS) and zeta potential (Anton Parr). Drug loading was determined using Quant-it Ribogreen kits (Thermofisher) which lysed the nanoparticles to determine input loading and encapsulation efficiency.

### Flow cytometry

Tumors were digested in Liberase (0.4 mg/mL, Roche, IN, USA) and DNAse (0.2 mg/mL, Roche) and processed through 70μm nylon strainers. These single cell suspensions were plated and then blocked with CD16/CD32 Fc block (Biolegend, San Diego, CA, USA). Cells then underwent immunophenotyping with fluorescent antibodies (Supplementary Table 2) staining. For viability assessment, cells were counterstained with DAPI (BD Biosciences, Franklin Lakes, NJ, USA). All cells were read with the BD-LSR Fortessa and analyzed using FlowJo.

### Measurement of cytokine, chemokine and DAMPs

Blood was collected retro-orbitally in heparin tubes and plasma was separated from blood using centrifugation and stored at -80°C. Tumors were submerged in 5 mL PBS and homogenized with a tissue homogenizer and then stored in -80°C. Samples underwent 3 freeze-thaw cycles and were centrifuged before the final aliquot storage in -80°C. Multiplex immunoassay (LegendPlex, Biolegend) was used to assess CCL2, CXCL10, IL-10, IFNγ and TNFα following the manufacturer’s instructions and read with the BD-LSR Fortessa. ELISA was used to quantify HMGB1 (CusaBio, Houston, TX, USA) in tumor tissues following the manufacturer’s instructions and absorbance readings were measured using a plate reader (Biotek, Winooski, VA, USA).

### Statistics

Statistics were performed using GraphPad Prism and data is represented as mean ± SD. Unpaired t-tests and one-way analysis of variance (ANOVA) were performed with the appropriate post-hoc tests. Multiple comparisons were conducted by comparing each group to each other and p < 0.05 was considered significant. All significance, if not specified follows the GP standard.

## Acknowledgment

This work was supported by grants from the National Institute of Biomedical Imaging and Bioengineering (R01EB028144, AAE), National Cancer Institute (R01CA253627, R01CA278633, E.K.), and pilot funding from the Case Comprehensive Cancer Center Support Grant (P30CA043703, AAE and EK). A.B. was supported by a fellowship from the NIH Interdisciplinary Biomedical Imaging Training Program (T32EB007509). We acknowledge the Imaging Core, Cytometry and Imaging Microscopy Core, and Tissue Core at Case Western Reserve University. We also acknowledge Taylor Moon, Inga Hwang and Laolu Ogunnaike for their assistance in flow cytometry studies, and Roshnee Mrinalini Chatterjee for illustrating the graphical abstract. Figure illustrations were made using BioRender.

