## Supplementary Information for "Enhanced delivery of lipid nanoparticle-based immunotherapy by modulating the tumor tissue stiffness using ultrasound-activated nanobubbles"

**Contents**

**Table 1.** Formulation details of polyacrylamide phantoms
**Table 2.** Flow phenotyping for immune cell populations
**Figure S1.** SWE phantom validation
**Figure S2.** SWE qualitative measurements
**Figure S3.** Characterization of lipid nanoparticles
**Figure S4.** Immune signaling (IFNγ)

| Theoretical elastic modulus | Acrylamide (mL, wt.%) | Bis-acrylamide (mL, wt.%) | Solvent (mL) | APS (mL) | TEMED (μL) |
|---|---|---|---|---|---|
| **1 kPa** | (1.7, 6.4) | (0.5, 0.095) | 7.8 | 0.5 | 50 |
| **10 kPa** | (4.0, 15.2) | (0.5, 0.095) | 5.5 | 0.5 | 50 |
| **18 kPa** | (7.0, 26.5) | (0.5, 0.095) | 2.5 | 0.5 | 50 |

**Table 1.** Formulation details of polyacrylamide phantoms (10 mL batch)

| *Cell population* | *Immunophenotype* |
|---|---|
| *Pan-immune cells* | DAPI-/CD45+ |
| *Dendritic cells* | DAPI-/CD11b+/CD11c+ |
| *Macrophages* | DAPI-/CD11b+/F4-80+ |
| *mMDSCs* | DAPI-/CD11b+/Ly6chiLy6G- |
| *T cells* | DAPI-/CD3e+/CD4+ or CD8+ |
| *NK cells* | Live/CD3e-/NK1.1 |

**Table 2.** Flow phenotyping for immune cell populations

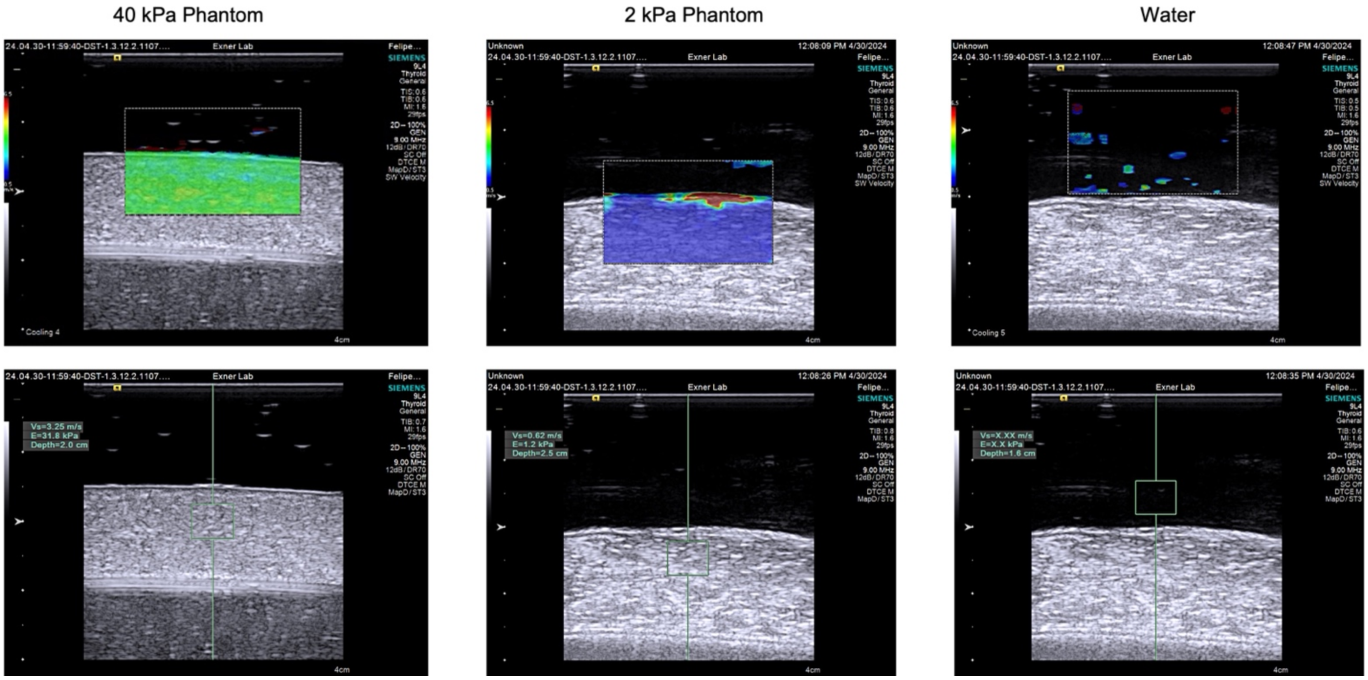


**Figure S1. SWE phantom validation.** Polyacrylamide phantoms were made based on the measurements in Table 1. Phantoms were placed in water and imaged using the Siemens 2000 using the 9L4 probe. Virtual Touch Imaging and Quantification Mode was used. The top row shows the qualitative elasticity map, while the bottom row displays the quantitative elasticity values measured within the region of interest (small box)

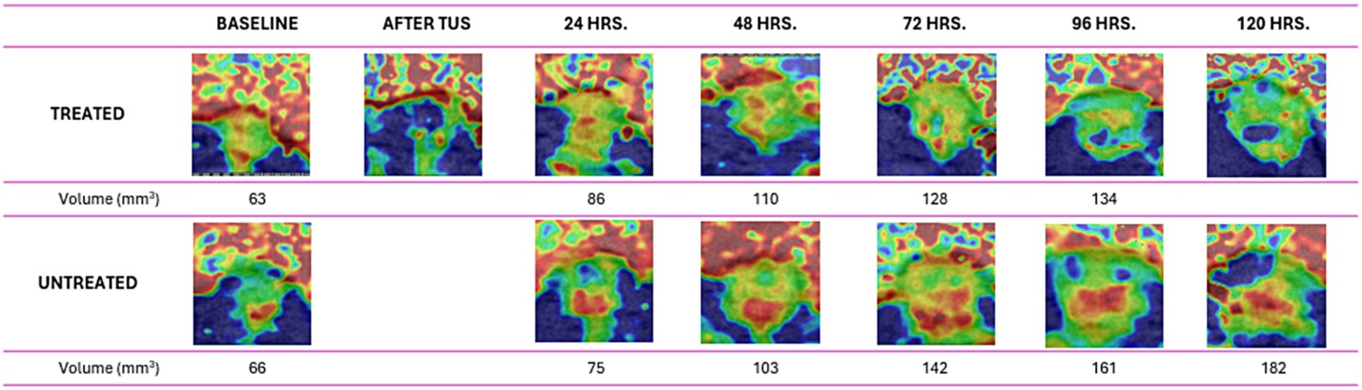


**Figure S2. SWE qualitative measurements.** (a) Uncropped elastograms for qualitative measurements. Due to the high spatial stiffness of the standoff gel pads used for the SWE setup, the elastograms were cropped to the tumor ROI and kept to their original size for ease of viewing of the growing tumor alone. Measurements taken by the Siemens 2000 using the 9L4 probe. Measurements are taken daily for 5 days. Set up was kept constant by regulating depth, gel thickness and probe angle. Qualitative elastograms are reported with the range of soft (blue) to hard tissue (red).

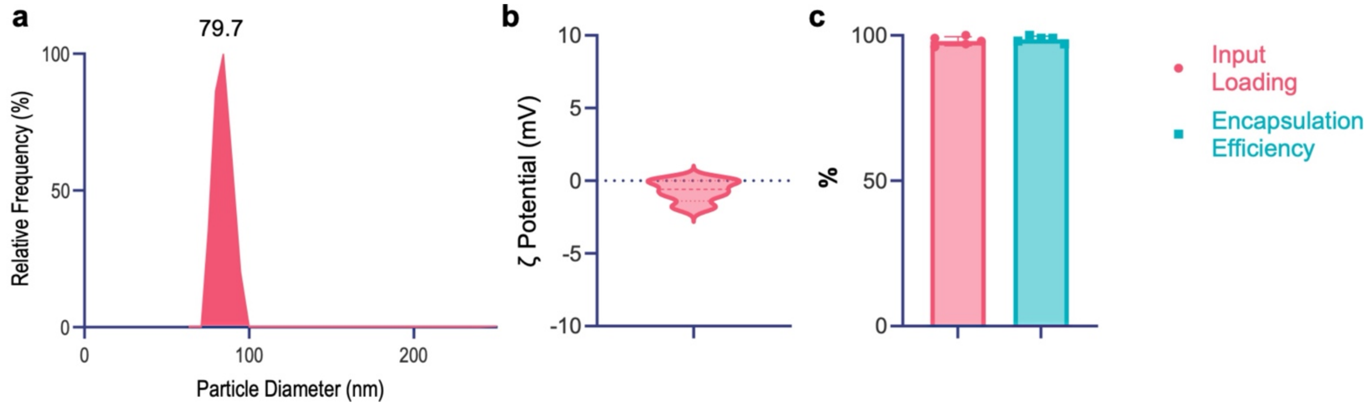


**Figure S3. Characterization of lipid nanoparticles.** Lipid nanoparticles carrying were formulated with the sonication method and characterized after 16-18 hours of dialysis. (a) Dynamic light scattering measurements were taken in PBS. The number-weighted distribution is visualized through a histogram to assess its dispersion. (b) Zeta potential measurements for surface charge evaluation are taken in DI water at RT averaged across a sample size of 3. (c) Drug input loading and encapsulation efficiency are taken by a fluorescent plate reader assay. Particles are lysed with 0.1% Triton X-100 at 37°C. Input loading is calculated considering the initial mass of drug added during formulation. Encapsulation efficiency is calculated by comparing lysed and un-lysed particles to determine the % of siRNA on the surface. N=3 was used for assessment.

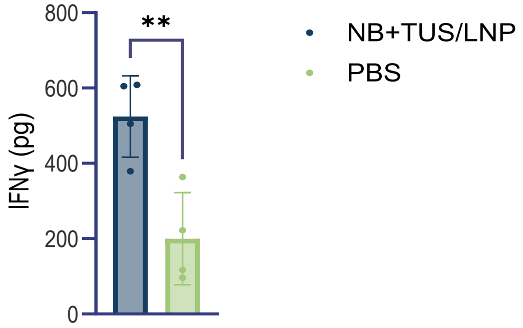


**Figure S4. Immune signaling for combination LNPs to assess long-term changes (IFNγ).** E0771.LMB tumors were treated on day 6,9,12 and harvested 24 hours after the third treatment. IFNγ was assessed for the NB+TUS/LNP and PBS groups using a multiplex bead assay. All concentrations were multiplied by the homogenization volume. All injection volumes across groups were kept constant. Statistics were carried out using a One-way ANOVA comparing the mean of each group to the others**.**
